# Effects of Parcellation and Threshold on Brain Connectivity Measures

**DOI:** 10.1101/522060

**Authors:** T.C. Lacy, P.A. Robinson

## Abstract

It is shown that the statistical properties of connections between regions of the brain and their dependence on coarse-graining and thresholding in published data can be reproduced by a simple distance-based physical connectivity model. This allows studies with differing parcellation and thresholding to be interrelated objectively, and for the results of future studies on more finely grained or differently thresholded networks to be predicted. The dependence of network measures on thresholding and parcellation implies that chosen brain regions can appear to form a small world network in many studies, even though the network of individual neurons may not be a small world network itself.

## 1. Introduction

It is important to understand the structure and function of the brain and its large network of 10^11^ neurons each with 10^4^ synapses to other neurons. In order to simplify this network to a manageable level, one of the main approaches used is to coarse-grain the network by grouping large numbers of neurons into regions (typically 30 – 1000, though some use 10000 – 30000) based on spatial proximity and then study the connections between these regions. The hope is that the properties of the coarse-grained network are similar to the original network of neurons, so that studying the former may provide useful insight into the latter; however, there is no proof that this will be the case. The connections between regions are usually determined by estimating the numbers of axonal fibers between them, but can also be based on functional correlations in their activity. From these measurements, either a connection strength is assigned between each pair of regions, or the network is thresholded to simplify the analysis, sometimes binarizing by keeping connections between regions only if they are above a certain strength and discarding them if they are weaker.

These regional networks are then typically characterized (Gong et al., 2009; He et al., 2007; Achard et al., 2006; Bullmore and Sporns, 2009) by using graph theory and calculating graph measures of the network, such as the clustering coefficient and the average path length. From this, conclusions about the structure of the overall network of neurons are often drawn, usually including a conclusion that the regions of the brain form a small-world network. This is done by comparing the network measures to those of the model in Watts and Strogatz (1998), which is widely used as a model for small-world networks. However, this analysis is done without directly acknowledging that the coarse-grained network of regions is an approximation to the physical network of the brain, and that information is also lost by binarizing connections. In addition, there is a large degree of variation in the properties reported between studies (Gong et al., 2009; Salvador et al., 2005; Iturria-Medina et al., 2008; He et al., 2007; Achard et al., 2006). There has been preliminary work on how the coarse-graining affects the network properties (Fornito et al., 2010), suggestions that the spatial embedding of regions will influence the path length and clustering (Nunez and Cutillo, 1995; Barthélemy, 2011), and work on spatial networks in general (Barthélemy, 2011, 2003), but no systematic analysis has yet been done the brain. Without ensuring that the analysis is robust with respect to changes in the parcellation and threshold, it is entirely possible that as the number of regions increases, the graph properties may change in ways that invalidate the analysis and its conclusions. For example, that a network of 100 brain regions forms a small-world network does not necessarily mean that their 10^11^ neurons form a small-world network too. For this conclusion to be drawn, a causal link would need to be found explicitly, instead of just being assumed to exist.

In addition, most network studies do not have any model for the process of obtaining connections between regions from the network of synapses when it is parcellated and thresholded. Many models have been put forward for the structure of the connections between regions [e.g. Vértes et al. (2012)], usually favoring preferential attachment [discussed in detail in Barthélemy (2011)], but sometimes following other rules. However, they are usually proposed ad hoc to match the data on coarse-grained connections between the regions, instead of modeling the underlying neuronal connections (which are ignored). To overcome this deficiency, we must start with a plausible physical model and coarse-grain it instead of jumping directly to modeling the regions.

In this paper we show that by simulating and examining the underlying network of individual neurons within the brain, we can predict how the coarse-grained network changes with parcellation and threshold. In particular, we predict how the clustering coefficient and average path length change with parcellation and threshold, and reproduce the data of Fornito et al. (2010). This analysis is then extended to determine how the small-world properties of the coarse-grained networks vary with parcellation and threshold.

## 2. Methods and Theory

In this section, we outline the procedure used for generating networks of regions from a simulated synaptic network.

In order to show all of the steps of the modeling procedure before applying them to real brain geometries, we illustrate them explicitly for a small network (36 regions of 5 neurons each), with the regions spread evenly on a 2D square (instead of approximately evenly throughout a 3D sphere, like an actual brain). This makes it easier to show how the synaptic network is thresholded to yield a regional network. Figure 1 shows the first step of placing the neurons throughout a simulated brain. Each of the 36 regions contains exactly 5 neurons, and there is some clumping of neurons occurring due to the random placement of them within the regions.

**Figure 1:**
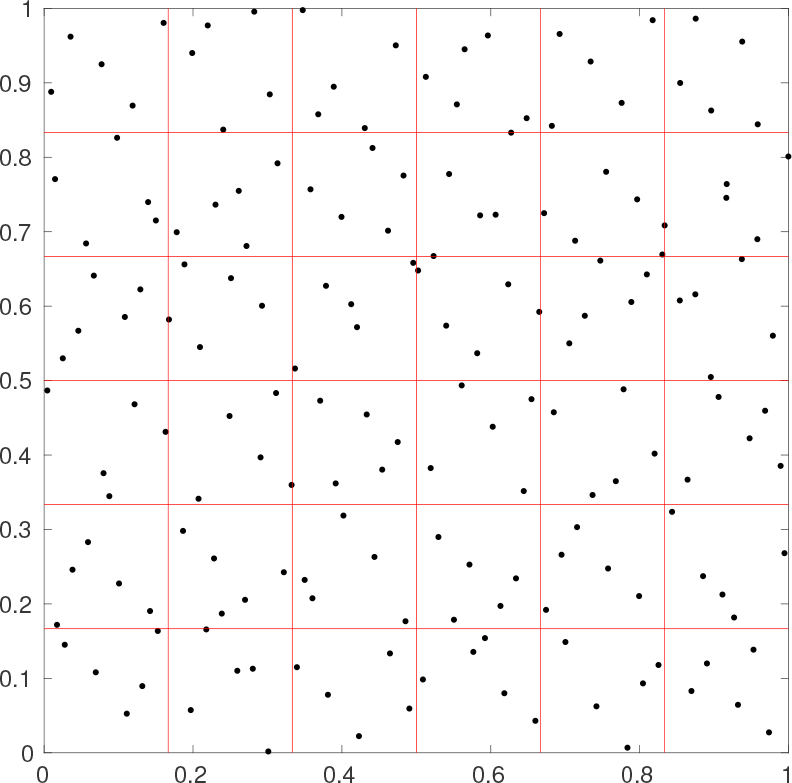
Locations of the neurons (black crosses) in the simulated network on a square cortex, in 36 regions of 5 neurons each (red lines show the borders of each region, axes are spatial coordinates on the square cortex with side length 1).

Once the neuron locations are determined, the next step is to determine connections between the neurons. For pairs of neurons, the probability of there being a synapse connecting them decreases the larger the further separated they are. Because longer range connections are rarer, possibly due to the cost of growing long axons, we assume the probability of connection between neurons follows a simple exponential distribution, although other distributions can be used. If the neurons are at positions **a** and **b**, the probability they are connected is approximated as

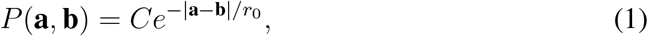

where *C* is a base connection probability between neurons located in very close proximity, and *r*_0_ is a typical connection range. The connections from neurons in one of the regions to all the other neurons can be seen in Fig. 2. This shows that the neurons are often connected to neurons within the same or adjacent regions, and rarely connected to neurons in distant regions.

**Figure 2:**
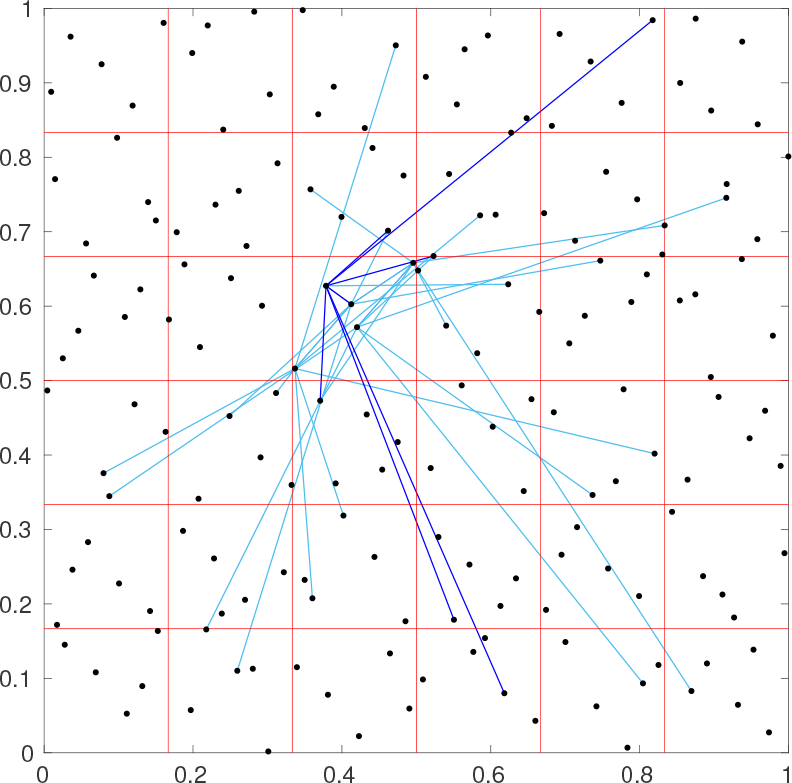
Connections (blue lines, darker for one particular neuron) from neurons in one region to ones in the rest of the simulated network, with the same neurons and regions as Fig. 1.

We now have a full simulated network of neurons and regions with a network of connections present between neurons. However, for a typical brain with 10^11^ neurons, this network is too hard to measure directly, so papers such as those by Gong et al. (2009); He et al. (2007); Achard et al. (2006) measured the connections between regions instead, by coarse-graining the network. The most common procedure is to examine the degree to which each pair of regions is connected (either through correlations of activity or the number of connections between neurons). Then, all connections between regions that are above a certain threshold are kept, and the rest are discarded. This gives a much simpler network that involves only connections between approximately 1000 regions instead of connections between all of the 10^11^ neurons.

The effects of thresholding can be seen in Fig. 3, where the number of interregional synaptic connections required for pairs of the 36 regions to be considered to be connected is set to either 1, 2, or 3, respectively. Each point in this figure corresponds to one of the red boxes in Fig. 1. The number of connections between the regions decreases with increasing threshold, and the majority of the connections are between regions in close proximity.

**Figure 3:**
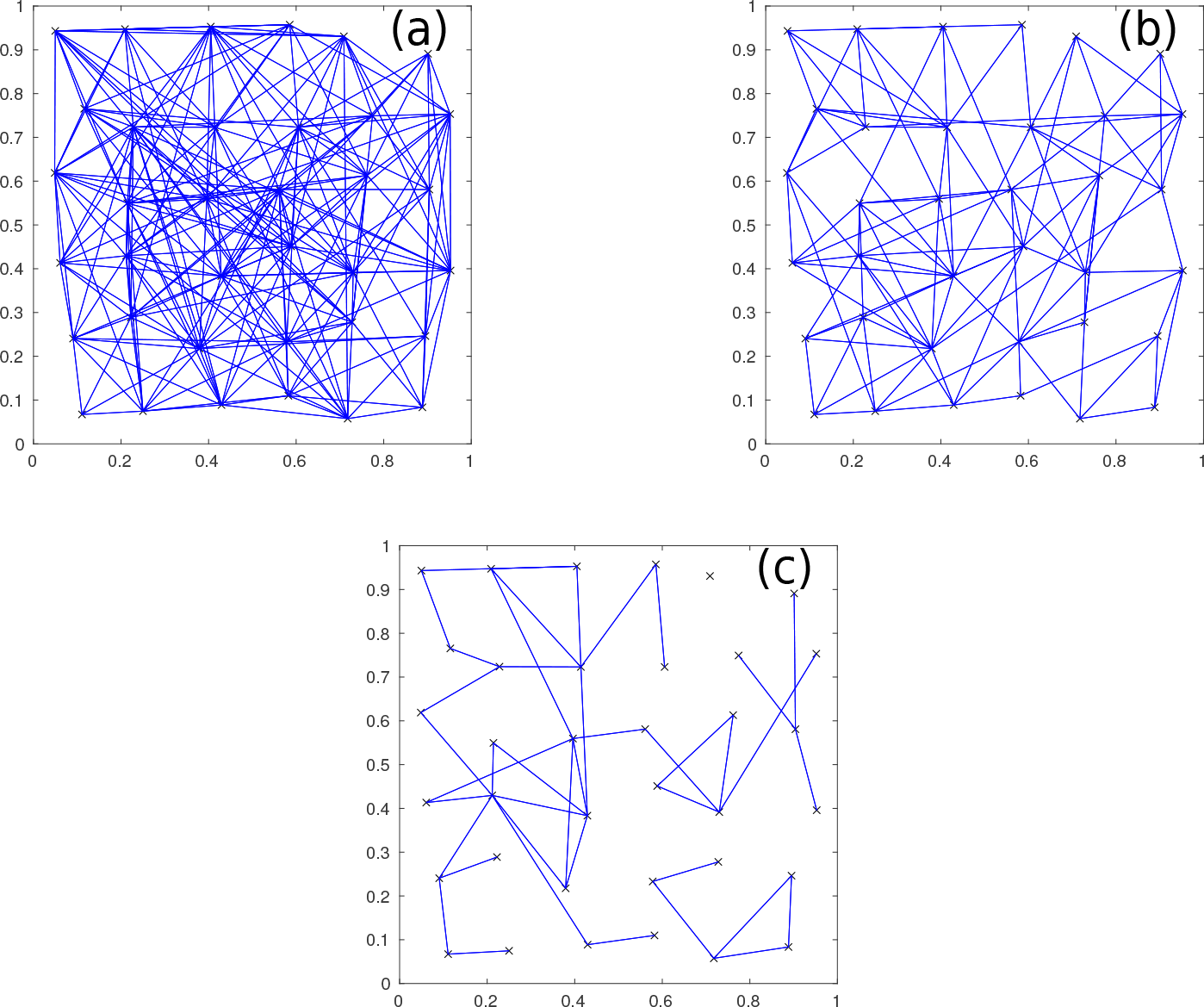
Connections remaining after thresholding between the 36 regions in Fig. 1 for three different thresholds [(a) is a threshold of 1 connection, (b) is 2, and (c) is 3].

To test the behavior for larger networks, we increase the number of neurons from 5 per region to 100, and compare to the previous results. This gives Fig. 4, where the threshold for the connections required is 200, 600, or 1000, respectively. This shows networks that are much more grid-like, because the correlation between proximity and connection strength between regions is much stronger. Decreasing the threshold gradually allows for longer connections between regions to be present. This behavior also tends to hold for other probability distributions for connections that decrease with distance, because the large number of neurons per region makes it very unlikely that distant regions will be strongly connected.

**Figure 4:**
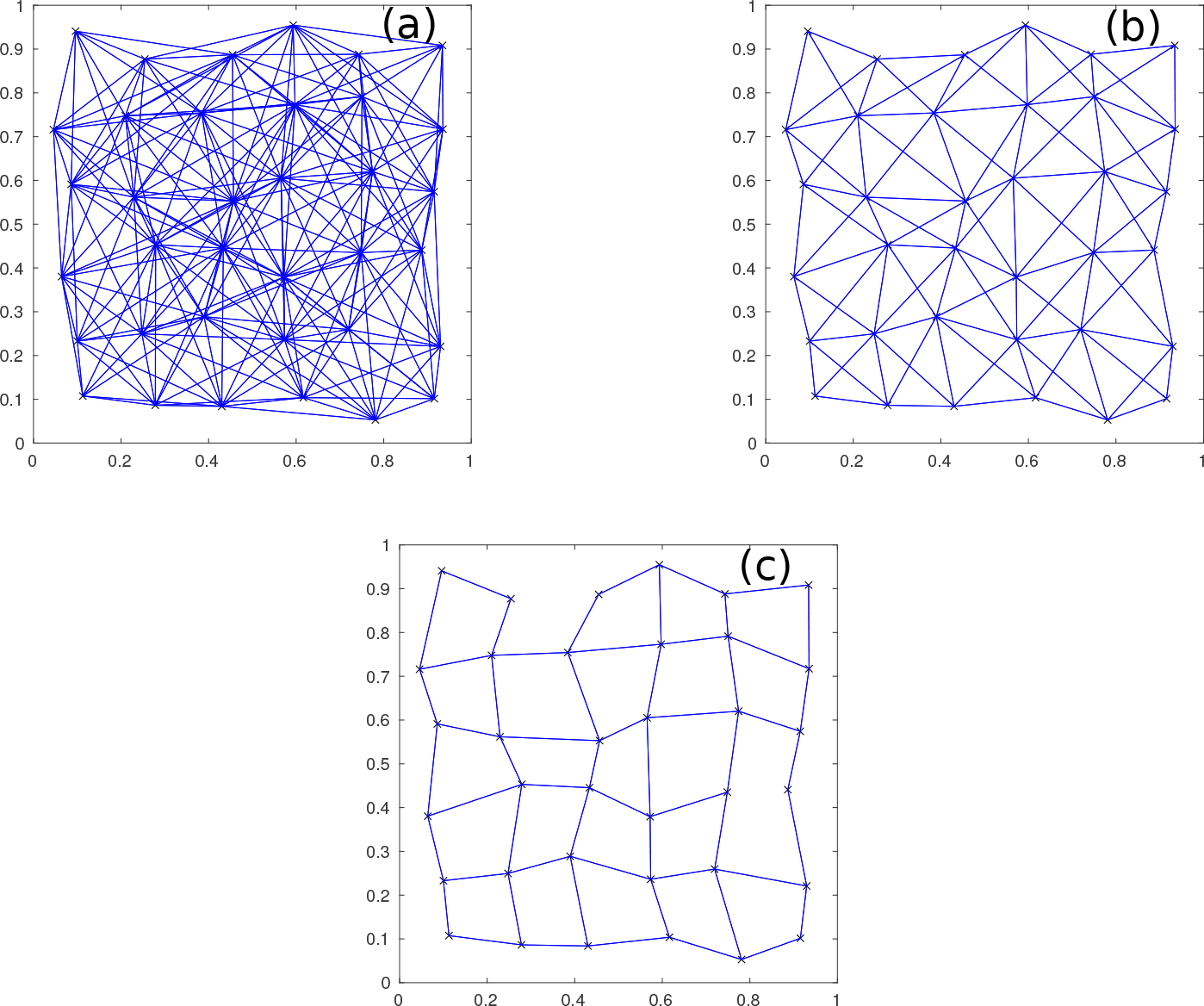
Connections between regions for 36 regions of 100 neurons, for three thresholds [(a) is a threshold of 200 connections, (b) is 600, and (c) is 1000].

The above behavior can be used to enable a very simple but effective model-based analysis of the coarse-graining. Because there will be approximately at least 10^7^ neurons present in regions at the parcellation scales used in experiments, thresholding will tend to just connect the closest pairs of regions together. This means that the coarse-graining of the neurons can then be modeled without having to simulate every neuron present.

Unlike the above example, which used square regions to cover a 2D plane, the model analyzed here distributed the centers of regions using a quasirandom sequence so they are approximately uniformly separated without much random clumping (Train, 2000) to fill a 3D sphere, before each of them is moved in a random direction up to 10% of the region’s radius, to avoid forming an almost perfectly regular grid. This gives them an approximately uniform spread throughout the sphere (which approximates the geometry of the brain or a hemisphere), while also allowing for some variation in how they are spread, as seen in experiments. Other geometries can easily be used, such as allocating the regions to cover the surface of a 3D sphere (which would match the surface connections more closely) or on a 2D disk, and these are used in the Results section to determine which geometry best reproduces the data of Fornito et al. (2010).

For the calculation of the clustering coefficient and average path length, we use the same procedures as in Fornito et al. (2010). In particular, if a region is disconnected from the largest connected component of the network, then a finite value for the path length has to be artificially assigned to it for the average path length even to be defined. Here and in Fornito et al. (2010), such nodes are defined to have an average path length from that node equal to the longest path length observed between a pair of nodes in the largest connected component of the rest of the network. However, because the real brain is a fully connected network, any *R* and *θ* values that lead to a disconnected network indicate that the resulting graph is too sparse to provide a valid representation, and so any conclusions derived when this is the case are highly questionable.

## 3. Results

In this section, we explain the main predictions made by the model that relate to coarse-graining neuronal networks, specifically relating to how network measures change with the number of regions and the threshold and what remains constant, and compare these predictions with published experimental data.

### 3.1. Clustering Coefficient

One of the most common measures used for determining how a network is structured is its clustering coefficient *C*, which is the probability of two neurons or regions being connected provided they are both connected to a particular third neuron or region. This has been measured for regional networks many times previously (Gong et al., 2009; He et al., 2007; Achard et al., 2006) and is usually used, along with the average path length, to argue that the regions of the brain form a small world network (a network with high clustering but low average path length). For example, Fornito et al. (2010) used resting-state functional magnetic resonance imaging (fMRI) to measure the interregional covariances of activity at 7 different parcellation scales (corresponding to *R* = 84, 91, 230, 438, 890, 1314, and 4320). By thresholding their connections at various covariance levels, networks of the interregional connections were obtained.

Motivated by this experiment, we examine how the clustering coefficient in the model’s networks changes with both threshold (relating to the proportion of connections remaining) and number of regions, as shown in Fig. 5. Each *C*-*θ* curve in this figure corresponds to a particular number of regions used in the simulation, averaged over 100 simulations because network details change with the locations of the regions, which are pseudo-randomly distributed. This shows that when *R* ≥ 1000, the clustering stops changing significantly with varying *R* for the range of thresholds used. However, if *R* is decreased, the clustering of higher density networks (*θ* > 0.2) stays approximately the same, while the clustering at lower densities drops significantly. This means that for studies with *θ* < 0.1, the clustering coefficient obtained from network analysis will depend strongly on *R*, and this needs to be taken into account when comparing between studies.

**Figure 5:**
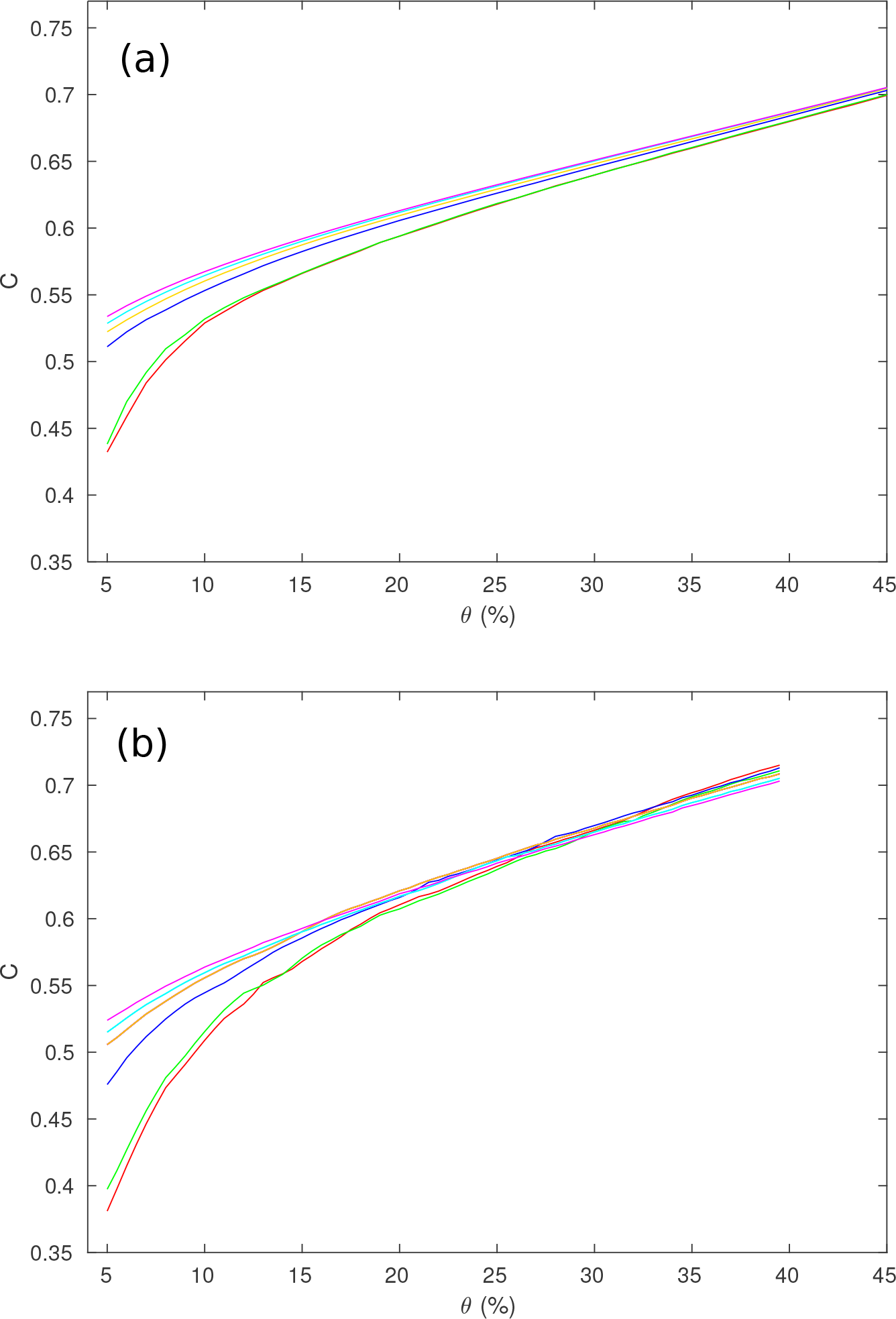
*C* vs. *θ* for different values of *R*, with *R* = 84 is red, 91 is green, 230 is blue, 438 is orange, 890 is cyan, 1314 is magenta. (a) Results from the simulated networks. (b) Results from the observed networks, adapted from Fornito et al. (2010).

We next compare the curves produced by the thresholding procedure with the data from Fornito et al. (2010), which showed the dependence of the clustering coefficient *C* on the threshold *θ* (called ‘Cost’ in their paper); their curves are shown in Fig. **??**. The main trends of the curves in both figures are the same, with the clustering being highest at high *θ*, and gradually decreasing as *θ* is decreased. Furthermore, the curves in Fig. **??** from the thresholded functional networks show the decrease in *C* for low *θ* as *R* decreases, as predicted by our model. The main discrepancy between the two figures is a slight disagreement in exactly how fast *C* decreases for very low *θ* values (the difference in values for *θ* > 0.15 is less than 0.02 for all parcellations) where the network is becoming disconnected and so the network analysis is not valid anyway, but the overall features of the plots are very similar in both figures.

For the previous simulation, we assumed that the regions were spread quasirandomly throughout a 3D sphere. However, it is possible that the regions are distributed differently, such as if they are concentrated on the surface, making the topology more like a 2D spherical shell. We test this hypothesis by adjusting the model to change how the regions are spread throughout the brain volume. Instead of obtaining the *C* curves seen in Fig. 5, we the obtain Fig. 6b if the regions are spread on a 2D disk, and Fig. 6c if the regions are spread on a spherical shell. As can be seen by comparing these plots to Fig. 5 and Fig. **??**, both the surface and disk topologies are significantly worse at predicting the experimental *C*, with both predicting much higher *C* at low *θ* for all *R* than Fig. **??**. This gives strong evidence that the changes in *C* seen in Fig. **??** are partially determined by the geometry of the brain regions as well as by *R* and *θ*.

**Figure 6:**
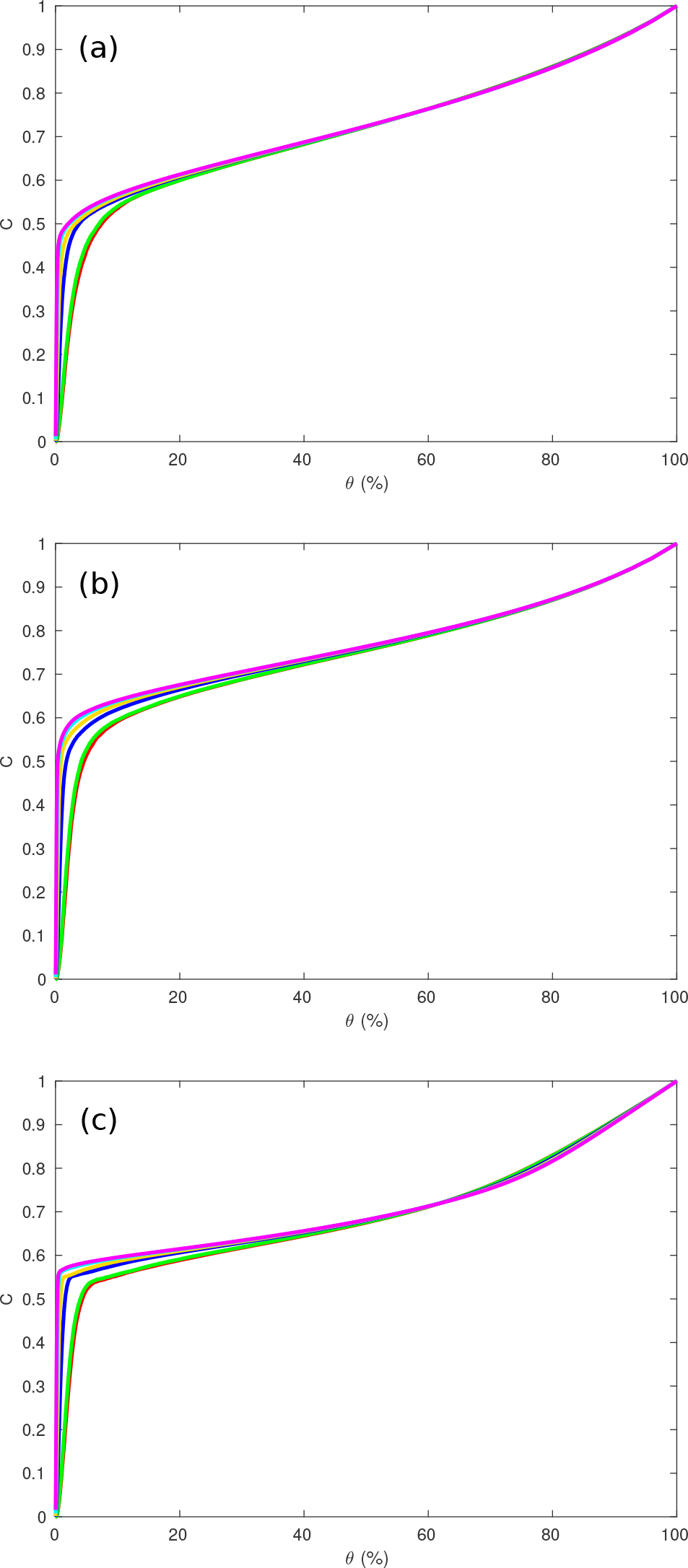
Same as Fig. 5 for the full range of thresholds, where lines correspond to the number of regions, *R* = 84, 91, 230, 438, 890, and 1314 from bottom to top. (a) Regions in a sphere. (b) Regions in a disk. (c) Regions on a spherical shell.

It is also important to note that any single simulated network may have *C* significantly higher or lower than that predicted by the above plots, which were averaged over many simulations. To get an idea of how much they vary, we simulate 400 networks and then instead of showing only the mean of *C* for each value of *θ*, plot the median, values separated by one standard deviation from the median, and values separated by two standard deviations from the median, which are shown in Fig. 7. This also shows *C* over the entire range of *θ*. This figure demonstrates that although the average of the clustering of the networks is highly constrained, any particular network can vary significantly. For example, only 90% of the networks simulated had *C* between 0.44 and 0.58 at *θ* = 10%. This further helps explain the variance of *C* observed in various studies, because not only will different values for *R* and *θ* significantly change *C*, but there will be variation beyond this, due to other random factors (such as the distribution of regions chosen). As an example, He et al. (2007) obtained *C* = 0.30 for their regional network, whereas Achard et al. (2006) reported *C* = 0.53 on a different network, which is much higher. However, their values for *R* were 54 and 90 respectively and their *θ* were 0.073 and 0.10. This explains the discrepancy, as *C* decreases in Fig. 5 as both *R* and *θ* decrease. This means that the discrepancies are consistent with being due to the different choices of *R* and *θ*, as opposed to actually having detected significantly different networks.

**Figure 7:**
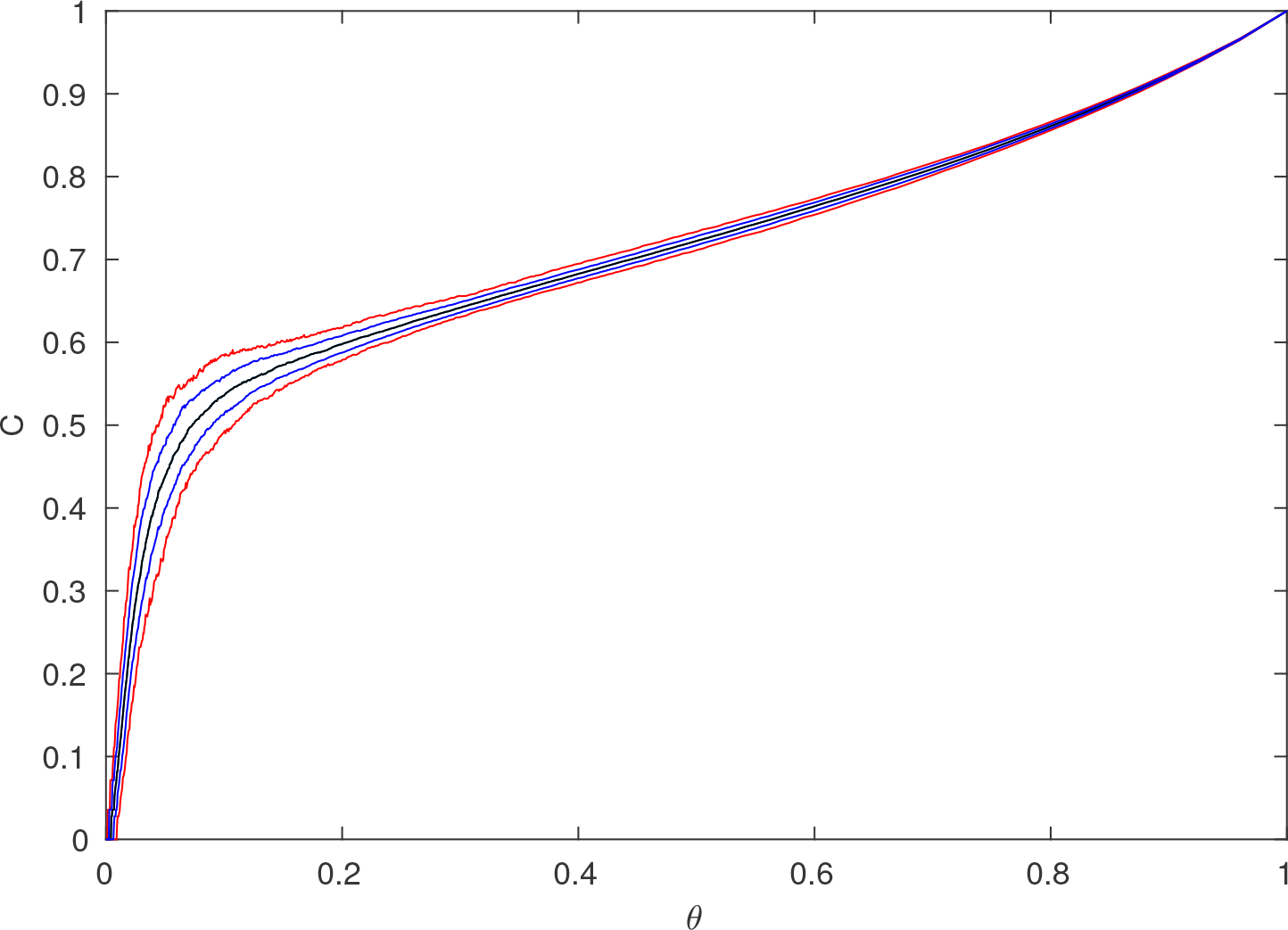
The median (black), one standard deviation from median (blue) and two standard deviations from median (red) clustering coefficients vs. threshold from 400 simulated regional networks, with *R* = 84.

We can explore these changes in *C* with *R* and *θ* further by simulating networks with intermediate values of *R* as well, and then examining the full 2D surface of *C* values, which can be seen in Fig. 8. With log scales on both axes, it shows that for *θ* > 0.1, the clustering does not vary much with *R*, which is supported by Fig. 5. However, as *θ* continues to decrease, the network starts becoming disconnected, making the clustering decrease rapidly to 0. The points at which this begins to happen fall on a straight line in Fig. 8, implying that it occurs for a constant value of *Rθ*. This can be checked by examining Fig. 9, which shows the clustering vs. *R* and *Rθ*. *C* falls rapidly toward zero as *Rθ* decreases below ≈ 5. By checking the size of the largest connected component against *Rθ*, it can be confirmed that this is the point at which this component starts rapidly decreasing in size, as large numbers of regions become disconnected from the network and the analysis is no longer relevant to the (fully connected) brain. This all suggests that the different rates of decrease of *C* seen as *θ* decreases in Fig. 5 are primarily due to the cases with lower *R* values reaching the *Rθ* = 5 threshold faster as *θ* decreases.

To compare these predictions of the model with experimental data other than the data from Fornito et al. (2010), we can overlay some of the *R*, *θ* and *C* values obtained in other experiments over Fig. 8, which are the black dots. In order of decreasing *R*, these are *R* = 90, *θ* = 0.1, *Rθ* = 9, *C* = 0.53 from Achard et al. (2006), *R* = 78, *θ* = 0.11, *Rθ* = 8.58, *C* = 0.49 from Gong et al. (2009), and *R* = 54, *θ* = 0.073, *Rθ* = 3.94, *C* = 0.30 from He et al. (2007). The model’s values for *C* at these points are 0.54, 0.54, and 0.44 respectively. This is a good agreement overall, the higher two values are reasonably close, while the discordant, lower value from He et al. (2007) is, from the data obtained, below the *Rθ* = 5 boundary, as shown in Fig. 9, indicating that the resulting network is significantly disconnected anyway, making any conclusions drawn unreliable. Even for the other two studies, the *Rθ* value is not that much higher than the boundary, due to the relatively small *R* used, demonstrating the general need for larger networks for these kinds of studies.

**Figure 8:**
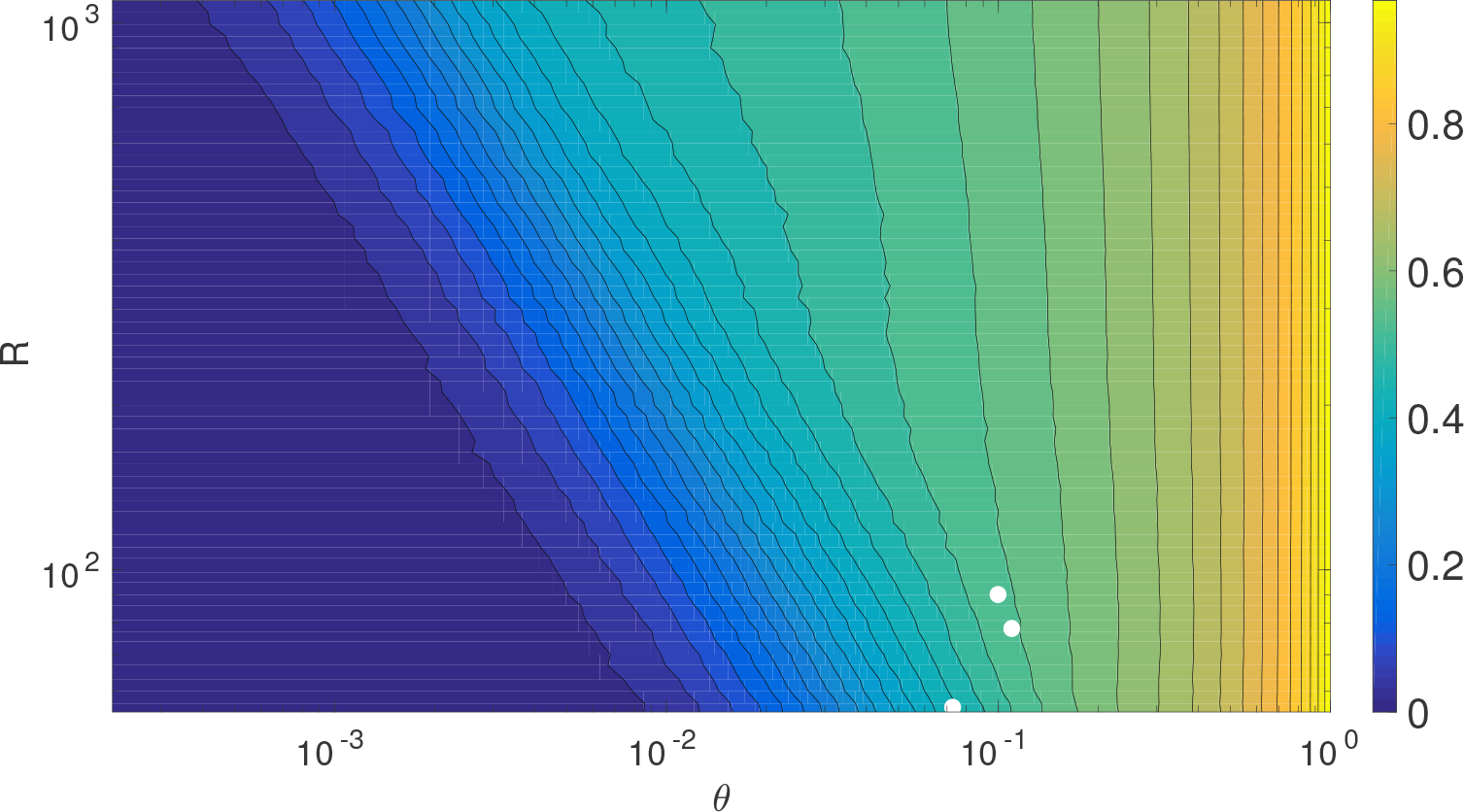
*C* vs. *R* and *θ*, with log scales on both axes and the color bar indicates the value of *C*. The dots show the *R* and *θ* values for three selected studies. In order of decreasing *R* (top to bottom), they are from Achard et al. (2006), Gong et al. (2009) and He et al. (2007).

**Figure 9:**
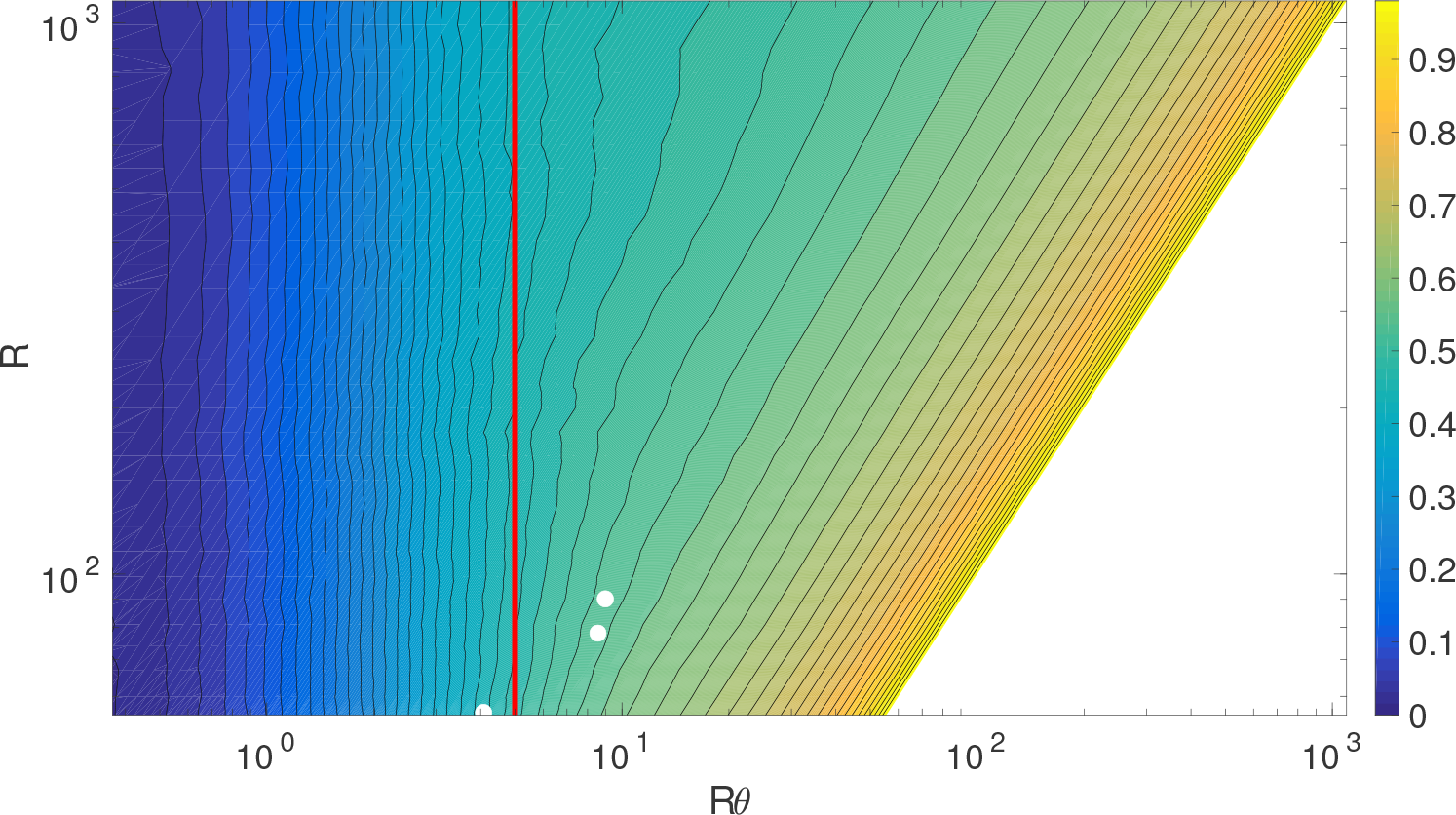
*C* vs. *R* and *Rθ*, with log scales on both axes and the color bar indicates the value of *C*. The dots show the *R* and *Rθ* values for three selected studies, and the vertical red line is *Rθ* = 5. In order of decreasing *R* (top to bottom), they are from Achard et al. (2006), Gong et al. (2009) and He et al. (2007).

In addition, Fig. 8 and Fig. 9 allow us to model much higher *R* networks accurately for *θ* > 0.1, because the *C*-*θ* curves approach a limiting curve quickly as *R* increases for that range, and for *Rθ* < 5, because this is consistently the point at which the network becomes disconnected and the analysis is no longer applicable to real brains. This can be applied to compare the model with van den Heuvel et al. (2008), which has *R* = 10000, significantly higher than the largest network simulated here. Their Fig. 2(c) shows *C* ≈ 0.50 ± 0.03 when *θ* = 0.15, decreasing to *C* ≈ 0.44 ± 0.02 when *θ* = 0.02. Estimating *C* from Fig. 8 by extrapolating the trends to higher *R* gives *C* ≈ 0.59 for *θ* = 0.15 and *C* ≈ 0.49 for *θ* = 0.02, overestimating *C* by a small amount in each case. Improving the model to account for the actual structure of the brain as opposed to the very simplified one used here would allow for very accurate extrapolation to large *R* networks.

### 3.2. Path Length

Another widely used measure for analyzing brain networks is the average path length between regions, which is the average number of connections required to join a random pair of regions in the network, and this is the other quantity usually used for determining small-worldness of a network. We can modify the above analysis to examine how the model predicts the dependence of average path length *L* on *θ* and *R*. Fig. **??** shows *L* vs. *θ* for the same values of *R* as in Fig. 5. This shows that *L* increases as *θ* decreases, with it increasing faster with *θ* for smaller *R*. This is as expected, because as *θ* decreases, the number of connections available to link a random pair of regions decreases, and so it will take on average more steps to link them to each other.

**Figure 10:**
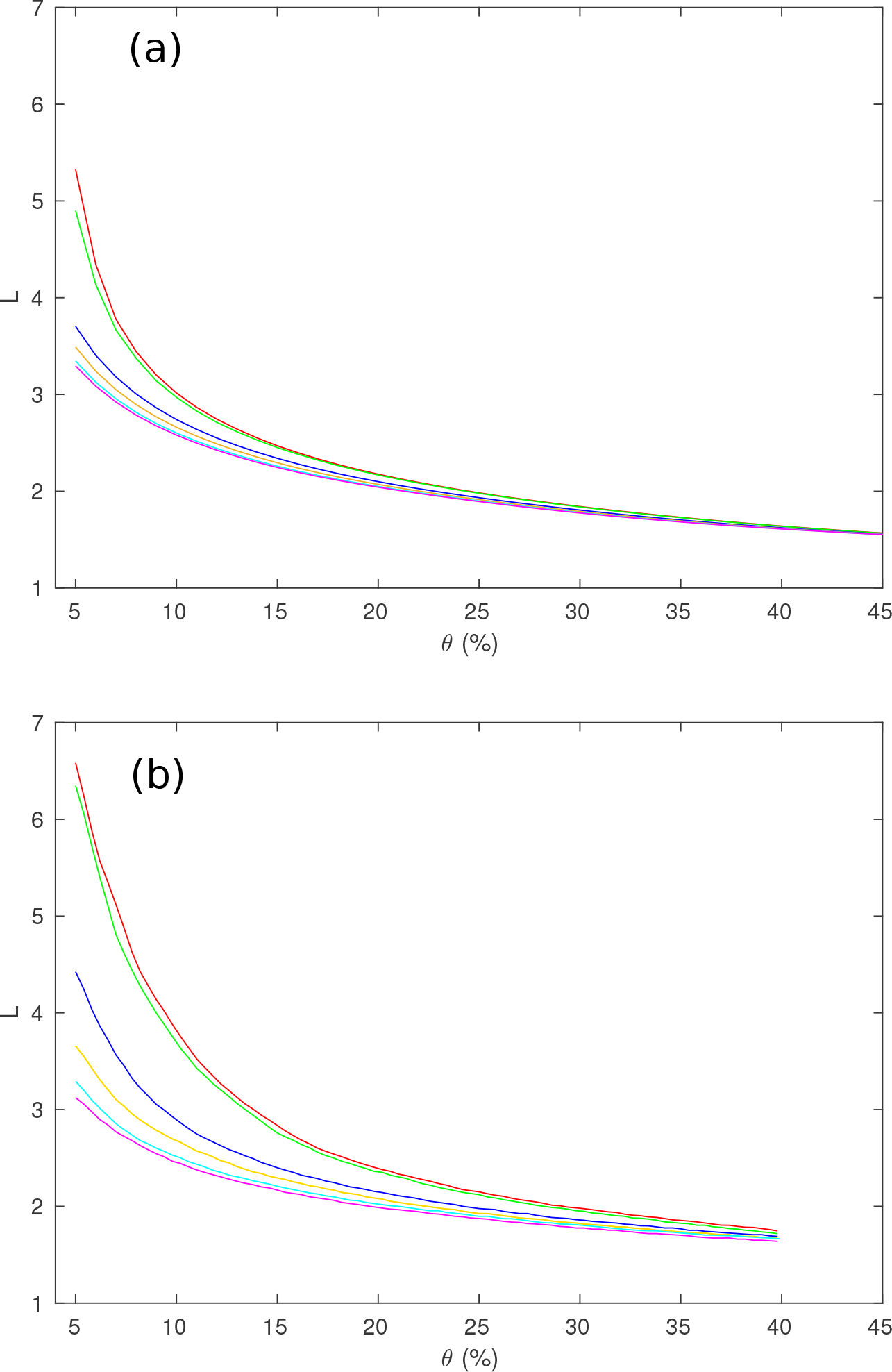
*L* vs. *θ* for different values of *R* (*R* = 84 is red, 91 is green, 230 is blue, 438 is orange, 890 is cyan, 1314 is magenta). (a) Results from the simulated networks. (b) Results from the networks adapted from Fornito et al. (2010).

We next compare the simulated *L* with the values found experimentally by Fornito et al. (2010), which are shown in Fig. **??**. The results are seen to agree very well with regards to the rapid increase as *θ* decreases (this comparison is done in more detail below). In both cases, *L* is minimal at high *θ*, and increases as *θ* decreases. Furthermore, in both cases, decreasing *R* gives a much higher *L*, due to regions becoming disconnected and the parcellation failing. The main difference is that the highest *L* seen in the model for *R* = 84 is only about 5.3 while in Fig. **??** it reaches 6.6 at *θ* = 0.05. The discrepancy is due to the model disagreeing with the experimental network over the size of the largest connected component of the network and artificial contributions from disconnected components. For such *θ* values, any conclusions drawn are unreliable, because trying to determine the properties of the completely connected brain by examining a coarse-grained network which is not completely connected is invalid from the outset.

Extending the range of *θ*, similarly to Fig. 6, gives Fig. 11. This clearly shows that *L* → 1 as *θ* → 1, and also demonstrates that *L* dramatically increases as *θ* → 0. Low *θ* values also demonstrate the behavior of *L* for networks that become disconnected (i.e. not real brain networks), because as the size of the largest connected component approaches 0, *L* → 0 from the artificial definition in this case.

**Figure 11:**
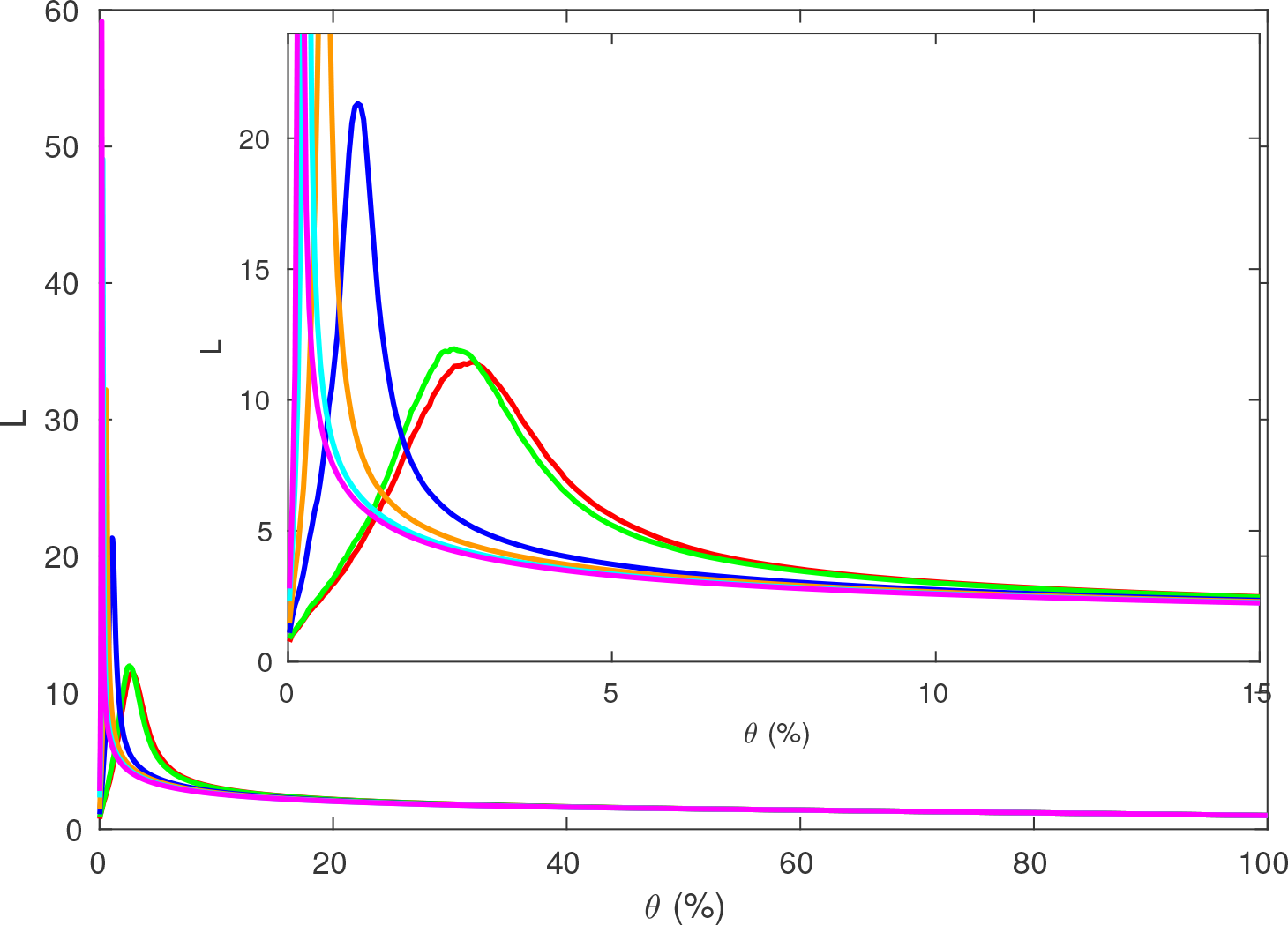
*L* vs *θ* as in Fig. **??**, but extended to cover all *θ*. The inset shows a closeup for low *θ* and *L*.

We now use the model to explain the behavior of *L*. To a first approximation, if *R* is large and *θ* is small, then each region will be connected to all regions with a small sphere around it, with a volume approximately equal to *θ* times the volume of the brain (because *θ* is the proportion of connections present). Hence, because the average distance between randomly selected points within the brain is a constant, *L* should be proportional to the average distance between two randomly chosen regions divided by the average distance that can be covered each step (*D*_*step*_). Because we have

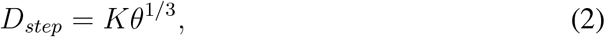

for a constant *K* (because the volume is proportional to the cube of the radius), *L*
should obey

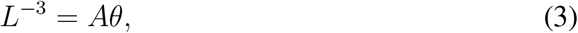

for a constant *A*. From the average separation of a random pair of points within a sphere, which is 36/35 times the radius (Burgstaller and Pillichshammer, 2009) and Eq. (3), we have

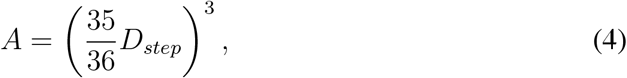

where *D*_*step*_ is the average step size between regions, as a fraction of the largest possible step (within the sphere of radius *θ*^1/3^).

In Fig. 12 we show this prediction for the networks used in Fig. **??** by plotting L^−3^ against *θ*, while also testing other geometries as in Fig. 6. This shows a clear linear relationship between these two quantities for a spherical geometry (for *θ* > 0.1, above the level below which the network becomes disconnected and the thresholding becomes invalid), with the line shifting upwards (i.e., A increases) and approaching exact proportionality with increasing *R*. When the network starts becoming disconnected at low *θ*, the path length drops suddenly, so *L*^−3^ diverges. The relationships for the disk and spherical shell geometries differ significantly from the sphere, and are clearly not linear over the range of thresholds shown.

**Figure 12:**
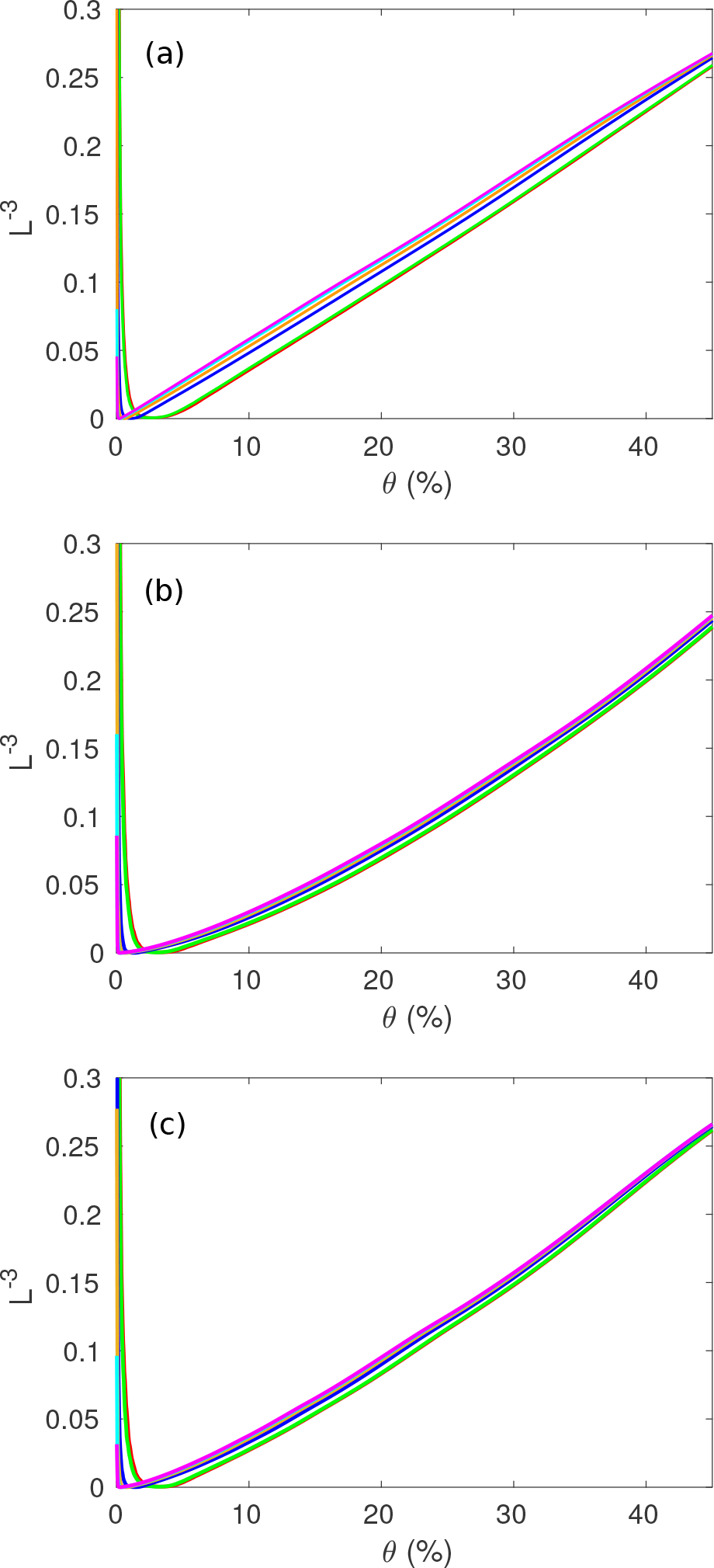
*L*^−3^ in the simulated regional network (averaged over 100 networks for each curve) vs. *θ* for changing *R* where *R* = 84 is red, 91 is green, 230 is blue, 438 is orange, 890 is cyan, 1314 is magenta. (a) Regions in a sphere. (b) Regions on a disk. (c) Regions on a spherical shell.

We compare the simulated curves with data from Fornito et al. (2010) in Fig. 13. While the lines are slightly shifted upwards compared to Fig. 12a, the experimental data from Fornito et al. (2010) clearly exhibit the same relationship as the spherical geometry, and significantly differ from the behavior for disk and spherical shell geometries. This strongly indicates that the behavior of *L* vs. *R* and *θ* in Fornito et al. (2010) is mainly due to how the regions are arranged in space, reinforcing the conclusion from the behavior of *C* as well.

**Figure 13:**
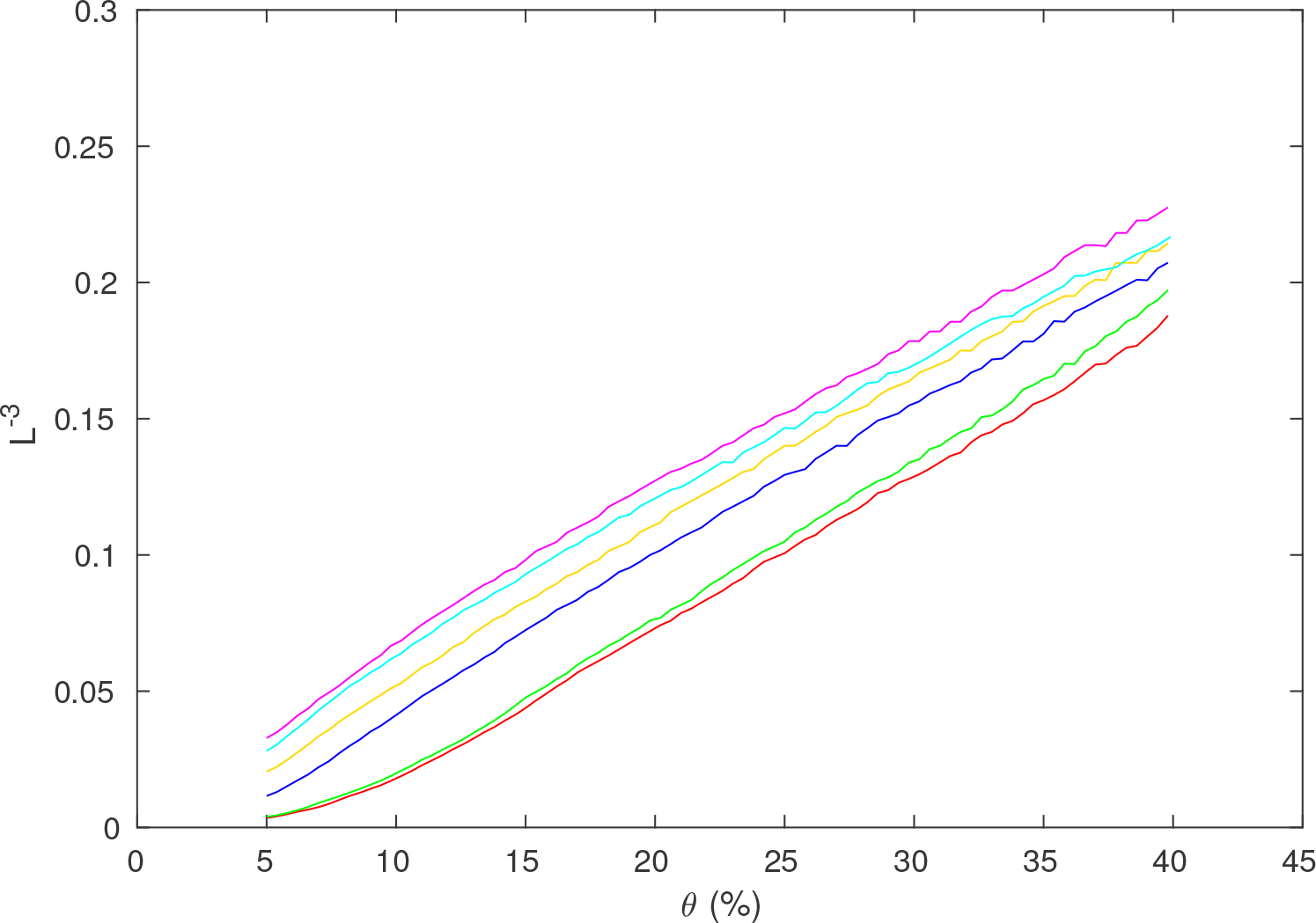
Dependence of *L*^−3^ vs. *θ* for same data as in Fig. **??** for different values of *R*, *R* = 84 is red, 91 is green, 230 is blue, 438 is orange, 890 is cyan, 1314 is magenta.

Finally, we estimate how closely the measured and predicted values of *A* from Eq. (3) match. Taking the slope of the *R* = 1314 lines for both the model and the data give an approximate value of *A* of between 0.55 and 0.60. This means that for the model and data that *D*_*step*_ ≈ 0.86, where the value *D*_*step*_ < 1 is due to connections between distant regions not being able to travel along the straight line connecting them, due to other regions not being present exactly along this line. This also suggests that *D*_*step*_ as found here will change with *R*, as higher *R* results in higher numbers of points lying close to this line.

### 3.3. Small-Worldness

The original measure of whether a graph is small-world was

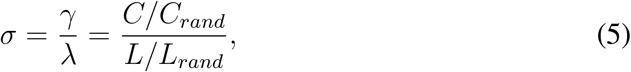

where C_*rand*_ and L_*rand*_ are respectively the clustering and average path length for a random graph of the same size and connection density (Watts and Strogatz, 1998). However, Telesford et al. (2011) argued that *σ* was overly sensitive to changes in C_*rand*_, and that larger networks tend to have larger *σ*, thereby making comparisons more difficult. Hence, they introduced an alternative measure *ω*,

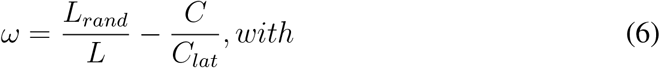

where C_*lat*_ is the clustering coefficient of an equivalent lattice graph, which is obtained by repeatedly swapping edges between nodes while preserving the node degrees until *C* reaches a maximum. Here, we will assume that the equivalent (same degree and number of nodes) lattice graph is topologically a 1D ring lattice, as discussed in Watts and Strogatz (1998), where the nodes are arranged in a ring and connected to their closest neighbors, which will be the case as long as all the regions have approximately the same degree. However, this is not spatially embedded, unlike the simulated brain networks.

We can examine how *σ* and *ω* change for the network of regions as *θ* varies, as shown in Figs. 14 and 15. Figure 14 shows that when less than 10% of connections are present, the value of *σ* is significantly above 1, indicating that in this region, the network will be considered to be small-world according to this measure. As *θ* → 0, *σ* dramatically increases to at least 20 in all parcellations, with *σ* > 600 for *R* = 1314. When the regions start disconnecting as *θ* approaches 0, *C* decreases faster than *L*, thereby decreasing *σ* in the process, but disconnection makes these thresholded parcellations invalid.

**Figure 14:**
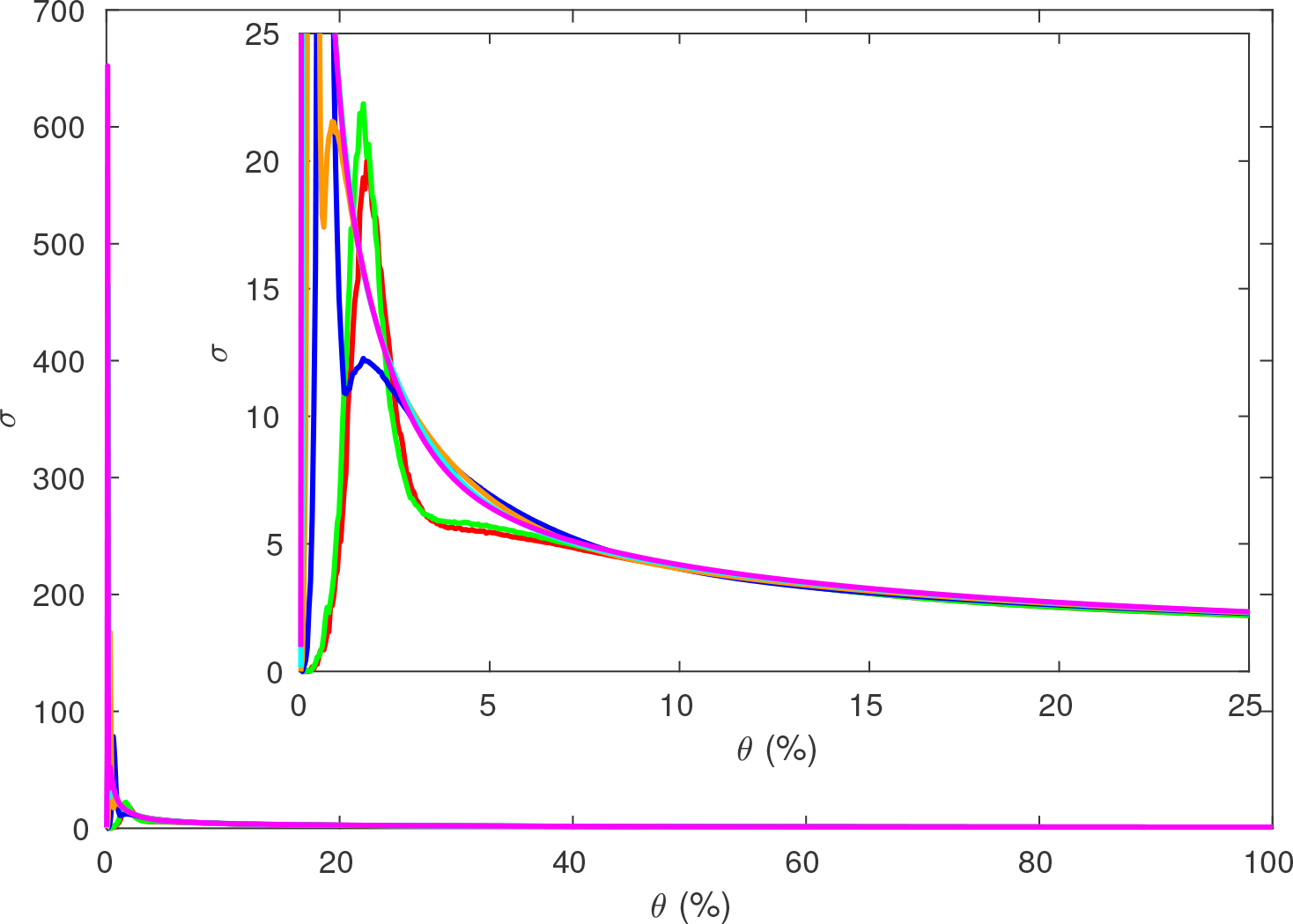
Small worldness *σ* vs. *θ* for the same data as Fig. 6, where *R* = 84 is red, 91 is green, 230 is blue, 438 is orange, 890 is cyan, 1314 is magenta. Inset is zoomed to show low *θ*.

**Figure 15:**
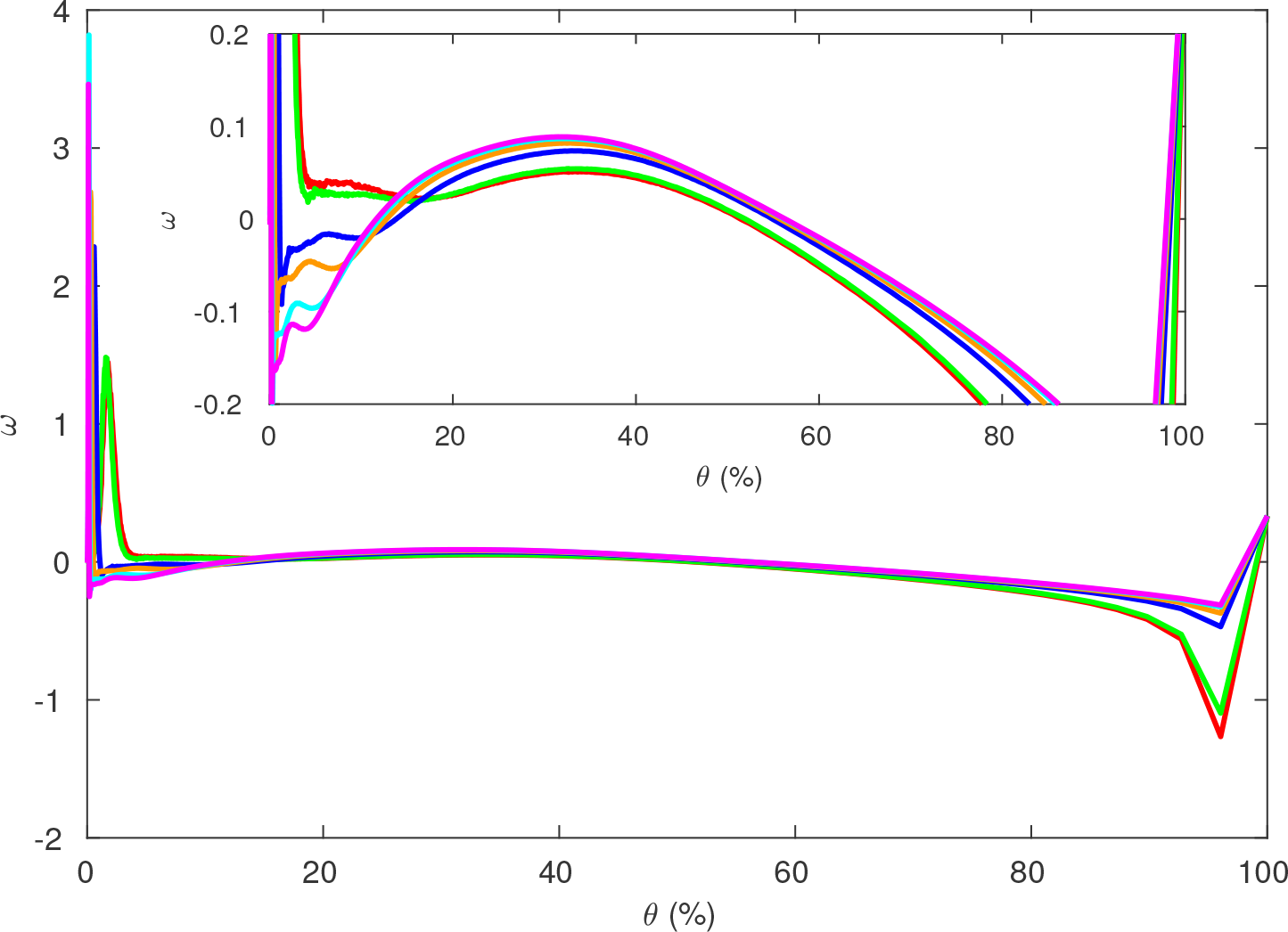
Small worldness *ω* vs *θ* for the same data as Fig. 6, where *R* = 84 is red, 91 is green, 230 is blue, 438 is orange, 890 is cyan, 1314 is magenta. Inset is zoomed to show low *θ*.)

Telesford et al. (2011) argued that |*ω*| ≤ 0.5 should be taken to indicate a small-world graph, with *ω* > 0.5 indicating a random graph, and *ω* < −0.5 indicating a lattice graph. Figure 15 shows that |*ω*| ≤ 0.1 for 0.05 < *θ* < 0.6, so that networks at these values for *R* and *θ* will always appear to be small-world graphs, and *θ* smaller than this leads to a disconnected network for most *R* and the results are thus not relevant to real brains.

However, the networks here are not small-world in the usual sense, because the simulated network of regions has all its connections between regions based on proximity. Only the closest pairs of regions are connected, and there are no long-range connections, which are usually considered to be required in small-world networks, to reduce *L*. The simulated networks’ misleadingly large *σ* and small |*ω*| values are at least partially due to *L* being much lower here than typical comparison networks, because the real network is embedded in 3*D* space. Hence *L* only increases as *R*^1/3^, in contrast to a typical lattice comparison network such as a ring lattice in which *L* ∝ *R*, making the brain network appear small-world due to the dimensionality of the network. This highlights the misleading inferences that can be obtained by comparing the brain to graphs that are not spatially embedded.

In addition, the small-worldness of the coarse-grained network of the brain doesn’t provide much information about the network structure at finer resolutions. When the thresholding is done, any long range neuronal connections end up being ignored because the much more numerous short range connections dominate the connectivity between regions, leading to important connections between distant regions being ignored. Hence, it is not possible to determine whether the neuronal network is small-world or not from studying the usual thresholded regional networks, because it is not possible to determine how many long range connections between neurons are present in the real brain by examining the thresholded network, from which they have been almost entirely deleted. However, Roberts et al. (2016) and Betzel et al. (2017) show that an alternate method is to keep the edges that are strong compared to how separated the regions are, not just the strongest overall. This preserves important long distance connections between regions, and helps mitigate the effects of the spatial embedding on the network measures.

## 4. Summary and Discussion

We have studied the dependence of graph properties of regional-level brain networks on parcellation scale *R*, threshold *θ*, and geometry by simulating the underlying neuronal network from a physical starting point and comparing with data. The main results are:

i. The simple physically based model produces networks of regions based solely on the distances between pairs of regions by assuming that neurons have a connection probability that exponentially decays with distance. When thresholded, this results in regions always connecting to their closest neighbors before any other regions. Despite the simplicity, the properties of these networks are very similar to the properties observed experimentally after thresholding experimental networks obtained via structural or functional measurements.
ii. The dependence of network properties such as the clustering coefficient *C* and average path length *L* on *R* and *θ* are accurately reproduced by the model, with only small discrepancies. This allows the results of different studies to be interrelated and discrepancies resolved, because many use only a single set of *R* and *θ* values and end up with different network properties. The ability to accurately simulate the network properties means that results for *R* and *θ* values other than those that have already been tested experimentally can be predicted, which is especially important for large *R* values, which are becoming increasingly relevant as experimental methods improve.
iii. The density of the final network can be freely set anywhere between completely connected and completely disconnected by varying *θ*. This means that information is lost by thresholding at any level and the information on variable connection strengths between regions should always be retained. However, this also means that new measures must be used in order to quantify connectivity, because some existing ones such as *L* do not generalize directly to variable connection strengths.
iv. If *Rθ* < 5, the regional network becomes significantly disconnected. Thus because the real brain is completely connected, all conclusions drawn in this region are unreliable and cannot be trusted because the network has been made too sparse to represent this fundamental aspect of the brain. Thus, care should be taken to not threshold any regional networks too strongly by choosing a very small *θ*, and the criterion *Rθ* > 5 must be tested and satisfied for any study to be valid.
v. The typical measures used to examine regional networks of the brain, including *C* and *L*, are not robust to changes in either *θ* or *R*, varying rapidly when either *R* or *θ* is small. This means that care needs to be taken when comparing the measures between different studies, to make sure that the variations due to *R* and *θ* are not changing the conclusions. Alternatively, they can use other measures that are more robust to changes than *C* or *L*, such as *L*^3^*θ*, so that these measures can be used for comparison between experiments instead.
vi. The measures *σ* and *ω* both indicate that the simulated networks are heavily small-world, despite not having any of the typical properties of a small-world network. The apparent small-worldness of these regional networks is mainly due to the spatial embedding of the regions in the brain into 3D space, because this embedding means that *L* ∝ *R*^1/3^, so the regions appear to form a small-world network even without any long range connections. Thus, when determining whether a network is small world or not, the reference networks it is compared to should be spatially embedded in the same way, so as to not let the spatial effects invalidate the comparison.
vii. The appropriateness of using thresholded networks to study the brain’s neuronal network is highly questionable. Conclusions about the network of regions, such as whether it is a small-world network, do not necessarily apply to the network of neurons, so conclusions about the architecture of the brain cannot be drawn from these measurements directly. Hence, when attempting to determine how the underlying neuronal network is structured, it is very important to understand how the structure of the neuronal network influences the thresholded regional network obtained. The network of regions should thus be seen as a simplified and transformed version of the network to be examined, not as the fundamental object of interest. However, methods such as those in Roberts et al. (2016) allow for a more robust method of thresholding which is not dominated by geometric effects, decreasing their influence on the results.

Overall the above results imply that standard thresholded network studies chiefly detect the properties of a near-neighbor network in 3D space.

## 5. Acknowledgments

This work was supported by the Australian Research Council Center of Excellence for Integrative Brain Function (ARC Grant CE140100007), by an Australian Research Council Discovery Grant (DP130100437), and by an Australian Research Council Laureate Fellowship (Grant FL140100025).

